# EpiBERTope: a sequence-based pre-trained BERT model improves linear and structural epitope prediction by learning long-distance protein interactions effectively

**DOI:** 10.1101/2022.02.27.481241

**Authors:** Minjun Park, Seung-woo Seo, Eunyoung Park, Jinhan Kim

## Abstract

**Motivation:** Epitopes are the immunogenic regions of antigen that are recognized by antibodies in a highly specific manner to trigger an immune response. Predicting such regions is extremely difficult yet contains profound implications for complex mechanisms of humoral immunogenicity.

**Results:** Here, we present a BERT-based epitope prediction model called EpiBERTope, a pre-trained model on the Swiss-Prot protein database, which can predict both linear and structural epitopes using protein sequences only. The model achieves an AUC of 0.922 and 0.667 for linear and structural epitope datasets respectively, outperforming all benchmark classification models including random forest, gradient boosting, naive Bayesian, and support vector machine models. In conclusion, EpiBERTope is a sequence-based model that captures content-based global interactions of antigen sequences, which will be transformative in epitope discovery with high specificity.

## 1 Introduction

The immune system is a complex defending system that defends and neutralizes infectious pathogens or harmful molecules [1]. The enhanced understanding of the immune system provides insight into immune-based therapeutics. At the molecular level, the immune system can be explained by the interaction of antigens and their corresponding receptors. For example, B-cells provide protection by producing antibodies, proteins that recognize and bind to unique pathogens (called antigens) in a highly selective manner to attack or neutralize the foreign substances invading the immune system [2]. These antigen amino acid (AA) residues that are recognized by antibodies are called epitopes. Therefore, identifying such epitopes is of great interest for the rational design of potential immune-based therapeutics and is an ideal starting point for drug discovery [3]. Accordingly, there have been many efforts to predict both linear and structural epitopes for the last decade. However, the performances of these methods have only been generally moderate [4]. One of the major challenges of the epitope prediction task is the limited size of the dataset. Exploring such vast antibody-antigen interaction domains with a small amount of dataset makes the prediction extremely difficult. In addition, predicting structural epitopes requires an understanding of the 3D folding structure of a protein, making the classification task even more challenging [5].

From a modeling perspective to choose relevant features, there have been two streams of applications. One is a structure-based prediction method that uses structure-related features [6]–[10]. A Structure-based method is an intuitive approach that utilizes 3-D structural information of the protein to identify epitopes that are closely located on the protein surface. This approach has a clear disadvantage that the model cannot train the antigen without a known structure. Furthermore, incorrect 3-D representation of antigens will increase the bias of the model. On the other hand, sequence-based prediction methods [11]–[13] only use amino acid sequences which are easily obtained by sequencing instruments. Hence, a sequenced-based method is cost-effective and time-efficient as it takes a primitive feature of protein sequences, yet it is less informative than a structure-based approach. To overcome this challenge, there have been efforts to use additional features such as hydrophobicity, polarity, predicted relative surface accessibility, and predicted secondary structure [11]. Their work improves the quality of predictions; however, some features come from another predictor [14] which stands as a nonreducible source of error. Furthermore, previous works included peptide sequences with a length less than 50 AAs taken from antigen sequences to predict the location of epitopes in a query peptide. This can lead to the loss of contextual information, limiting an understanding of the whole antigen sequence.

In order to address these challenges, we adopted Bidirectional Encoder Representations from Transformers (BERT) [15], an open-source transformer-based machine learning framework for natural language processing (NLP). Recently, the BERT architecture has gained popularity for utilizing the surrounding text of words to understand the context, also known as the attention mechanism. BERT has been highly successful in supervised tasks such as natural language inference [16] and machine translation [17]. The primary explanation for the outcome is that BERT is designed to read the sequence from both directions simultaneously, different from traditional methods of learning either left-to-right or right-to-left technique. Another noticeable characteristic of the BERT paper is how to train the model. Devlin et al. [15] use a large number of unlabeled text to pre-train the model by masking some tokens in the text and predicting masked tokens. Then, the pre-trained weights are served as initial parameters for other tasks such as question and answering, sentiment analysis, and sequence similarity classification. This subsequent procedure is called fine-tuning. During pre-training, the model learns how to make features from a large dataset. This feature is then used to solve downstream tasks which have relatively smaller datasets than the pre-train data. Pre-training and fine-tuning procedures are called transfer learning. Due to the flexible nature of transfer learning, pre-training becomes common in subsequent NLP research. Moreover, pre-training BERT on a large protein dataset [18] is widely studied and used to predict secondary structure, contact prediction, and remote homology [19]. Another work applying BERT on protein focuses on self-attention values which are intermediate features of the pre-trained model [20]. Self-attention can be interpreted as discovering relationships between tokens in a given sequence. Vig et al. visualize how self-attention correlates with contact map, binding sites, and substitution matrix and conclude that pre-trained BERT can learn those features solely from the sequence information.

In the application of the epitope prediction problem, EpiBERTope utilizes BERT architecture to learn antigen sequences from both directions of entire sequences, which aligns with the hypothesis that positive and negative epitopes are potentially affected by the entire amino acids in a sequence. In other words, EpiBERtope learns context-level relationships among amino acids within antigen sequences, which resolves the long-standing challenge of linking long-term dependencies in sequential problems. Instead of adding more features, EpiBERTope only takes amino acid sequences, pre-trained on a large protein dataset to learn embeddings of amino acids which can be effective while fine-tuning on the small epitope dataset.

As proof of concept, pre-trained and fine-tuned EpiBERTope is compared to traditional supervised learning algorithms, which have limited capability to interpret the entire context of antigen sequences. As expected, with a much broader knowledge of protein sequence grammar and global dependencies, EpiBERtope outperforms all benchmark classification models as well as published methods in all tasks. We also validated the significance of the pre-training data by comparing the model performance for each pre-training dataset: Swiss-Prot [21], TAPE [19], [21], and without-pre-training. Overall, EpiBERTope was able to attain high performance as a sequence-based model. Thus we believe that EpiBERTope’s transformer-based approach will have a significant impact on future studies to discover highly specific antibody-drug research.

## 2 Methods

### 2.1 Datasets

#### 2.1.1 Swiss-Prot database

EpiBERTope is pre-trained on the Swiss-Prot database [22]. Swiss-Prot is a manually curated database of UniProtKB that provides a high-level annotation of protein sequences, achieving minimal sequence redundancy and a high level of integration with other databases. This database contains more than 560,000 protein functions, domain structure, post-translational modifications, and identified variants. In the pretraining step, we extracted all sequences from the database and passed them to the model to find feature embeddings including the functional grammar of protein sequences. While pretraining, we randomly split the dataset to 6:2:2 ratio as train, valid, and test set.

#### 2.1.2 Linear epitope dataset

Linear epitope data is cataloged in the Immune Epitope Database (IEDB), where antibody and T cell epitopes are assayed in human and non-human species when exposed to infectious disease, autoimmunity, etc. [23]. We used the preprocessed dataset by Jespersen et al., which consisted of 11,834 epitopes and 17,722 non-epitopes. Here peptides shorter than five or larger than 25 amino acids were removed [11]. Furthermore, the positive and negative datasets only include those peptides cross-validated in two or more separate experiments. For EpiBERTope benchmark prediction, the dataset was randomly split into 80% train for cross-validation and 20% test dataset.

#### 2.1.3 Structural epitope dataset

We obtained a structural epitope dataset processed by Jespersen et al. [11] A dataset was collected from the Protein Data Bank (PDB), consisting of 675 structures of antibody-antigen complexes. To prevent redundancies, complexes with more than 70% of identical antigen residues to any other complex are removed, resulting in a total of 160 structures. To ensure the quality of the data, the remaining complexes have a resolution below 3 Ångström (Å), and all antigen sequences have more than 60 natural amino acid residues. To define the test dataset, a clustering program called CD-HIT was used with a 70% sequence identity threshold, from which 5 clusters (6 antigen sequences total). In total, we used a positive dataset of 3542 epitopes and 36,785 non-epitopes for training and testing combined.

### 2.2 Data preprocessing

#### 2.2.1 EpiBERTope preprocessing

After pre-training using the Swiss-Prot database sequences, the model is fine-tuned on structural and linear epitope datasets separately. Since the pre-trained model can take sequences with the maximum length of 2048, if the sequence length is greater than 2048, we ensure that all epitopes are preserved in the sequence by positioning epitopes or non-epitopes in the center. Sequences with a length less than 2048 are padded with zeros to make sure all sequences have equal sizes. For each antigen sequence, EpiBERTope tokenizes each amino acid with a predefined vocabulary list, enabling the model to interpret numerically.

For the model output, epitope and non-epitope residues are labeled as 1 and 0, respectively. The rest of the unknown regions are labeled as −100 to ignore in the loss function update (Equation 7)). In the final evaluation, only pooled positive and negative residues are considered in the performance metric.

#### 2.2.1 Benchmark ML models preprocessing

To compare EpiBERTope performance with other ML models, we tried an extensive set of ML classifiers, including ensemble methods, generalized linear models, naive bayesian methods, discriminant analysis, bagging-based and boosting-based classifiers. For the model input, we processed the sequences by positioning epitope or non-epitope residue in the middle with 4 flanking amino acids, resulting in 9-mer amino-acid sequences. Since resulting sequences are categorical variables, they are vectorized by concatenating 9 one-hot-encoded vectors, each representing the unique position of the 9-mer sequence. The final vectorized matrix is used for the benchmark ML model inputs

### 2.3 Pre-training EpiBERTope prediction model

EpiBERTope is a transformer-based language model that can be split into two stages: pre-training and fine-tuning. Instead of using a successful pre-trained BERT model such as the TAPE [19], we pre-trained with a manually reviewed protein sequence database called Swiss-Prot. The goal of the pre-training is to explore the set of hundreds of thousands of expertly curated protein sequences, learning the protein sequence grammar and amino-acid composition without introducing any biological and chemical properties. Briefly, each input protein sequence is first truncated if the length is greater than 2048. After truncation, each sequence is first transformed as tokens: *S* = (*t*_1_, *t*_2_, …, *t_n_*). For each protein sequence, EpiBERTope randomly masks 15% of the sequence and uses context amino acids surrounding a masked token to predict which amino acid residues fill in the masked region: (*p*(*t_masked_*|*t_unmasked_*). We used BERT-base architecture [24] that has 12 encoder layers, 252 hidden units, 6 self-attention heads (Equation 3), which computes the pair-wise interactions between each input amino acid with the rest of the sequence, enabling richer interpretations of long-distance interactions. For each protein sequence, the model provides (*N, H*) shaped matrix where *N* is the length of sequence and *H* is the number of hidden units.

#### Self-Attention layer

BERT is a bi-directional model that learns the global dependencies of the entire sequence of amino acids at once. It does so by computing the attention score *α* of each amino acid pair *i*, *J* in a protein sequence.

Attention by definition is a weighted sum of values:

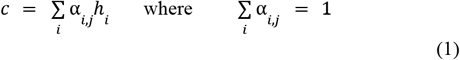

In Equation 1, *h_i_* represents a value vector, which decides how much information to use from other amino acids. To calculate the attention score *α_i,j_* we refer to transformer architecture [24], [25]. Briefly, the self-attention layer initializes three vectors: query (*Q*), key (*K*), and value (*V*). The attention score *α_i,j_* above is computed by projecting query vectors on key vectors by taking a normalized dot product of the two vectors (*QK^T^*). The output is converted into a probability vector:

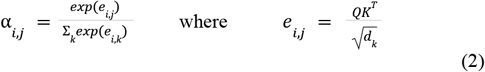

Finally, the model applies the weighted sum of the attention weights from value vector (*V*) to calculate the final attention score. The entire operations can be written as the following:

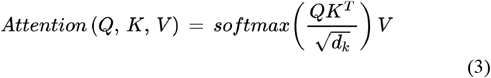

To capture a broader range of relationships between amino acids, the model learns 6 attention mechanisms (multi-head attention) that operate in parallel:

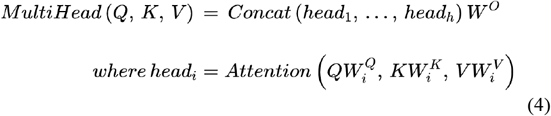

Here the projections are parameter matrices that are learned in pre-training. With a multi-head architecture, the model learns a deeper representation of the sequences, enabling richer interpretations of amino acid interactions. To prevent overfitting, validation loss was tracked every epoch, and training was stopped when the validation loss was not improved for 30 epochs. All hyperparameters for pre-training were chosen by grid-search with the lowest validation loss. We implemented the NLP model through HuggingFace Transformers [26]open-source platform based on *Pytorch [27]*.

### 2.4 Fine-tuning EpiBERTope prediction model

After pre-training with the Swiss-Prot database, EpiBERTope is fine-tuned on both the linear and structural epitope datasets. Instead of masking the inputs as in pre-training, the model performs a token classification of each residue (epitope or non-epitope) in the final layer. As described in Section 2.3, the model computes attention scores to learn long-distance interactions of antigen sequences, where residues per antigen sequence are connected with one another and the connection weights are dynamically calculated to understand global dependencies.

As shown in Figure 2, in the epitope prediction task, we represent the input antigen sequence as a tokenized input, introducing only one embedding. During fine-tuning, we only utilize a start vector *S* ∈ *R^H^* and an end vector *T* ∈ *R^H^*. In the final layer, the probability of the token *i* being an epitope is computed as a dot product between *T_i_* and *S* followed by a softmax over all residues in antigen sequence:

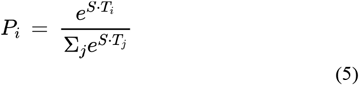

**Fig. 1.**
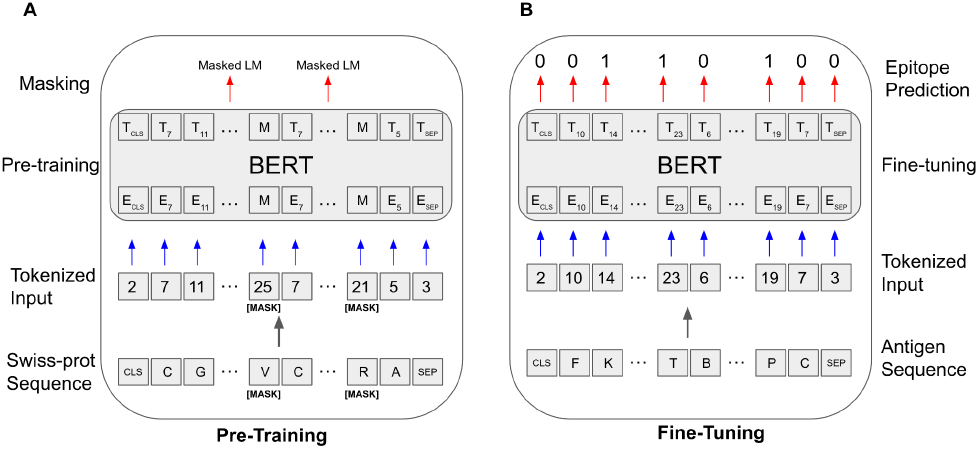
EpiBERTope model architecture. (A) Pre-training part of EpiBERTope. The model takes sequences from the Swiss-Prot database as inputs and randomly masks 15% of the sequence and predicts what amino acid should fill the masked regions. (B) Fine-tuning part of EpiBERTope. Input sequences consist of either linear or structural epitope datasets. Inside the BERT module, pre-trained protein sequence embeddings are utilized to classify epitopes and non-epitopes.

**Fig. 2.**
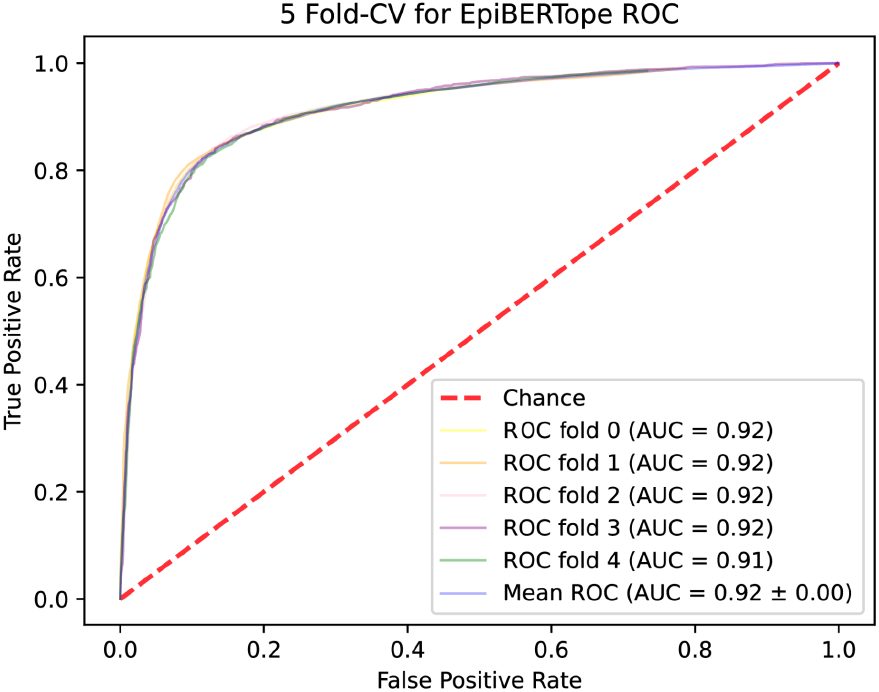
5-fold cross-validation ROC Curve. Optimal model hyperparameters are chosen based on the 5-fold cv result.

We actively searched for the best combinations of the learning rate, batch size, and warm-up step of the model. For training epochs, we chose the optimal point where the validation loss starts to increase.

### 2.5 EpiBERTope loss function

For the loss function, we used weighted cross-entropy loss. Specifically, we normalized the log-likelihood proportionally to the reverse of the initial weights so that the model assigns more weights on the label with smaller examples and vice-versa. This can be mathematically described as follows.

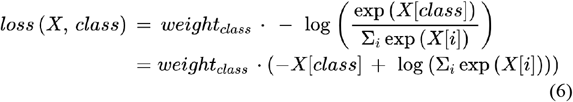

where:

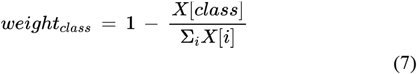

where the class can be either 0 for non-epitope or 1 for epitope amino acid.

### 2.6 ML models hyperparameter tuning

For hyperparameter tuning, 5-fold cross-validation is performed. In cross-validation, peptides originating from the same sequence are clustered together in either train or validation set, in order to prevent potential data leakage. In addition, the ML models are optimized through an exhaustive hyperparameter grid-search algorithm in *scikit-learn [28]*. The model selects the optimal parameters that maximize the score of the left-out data in cross-validation. The set of hyperparameters used for grid-search and the optimal combination can be found in Supplemental Table 1.

## 3 Results

### 3.1 EpiBERTope outperforms state-of-the-art results on diverse machine learning benchmarks

EpiBERTope is a trained transformer encoder stack that handles relationships between multiple sequences precisely from both directions of the sequences, leading to high performance on distinguishing epitopes from non-epitopes. As shown in the model architecture in figure 2, EpiBERTope learns protein sequence grammars in the pre-training step and then utilizes learned features to understand specific properties to distinguish epitopes and non-epitopes using antigen sequences alone.

For checking the stability of the model, we used a 5-fold cross-validation. For the linear epitope prediction task, due to the uniform size of the dataset, we observed a consistent and superior average AUC of 0.92 for the validation set (Fig. 2). Likewise, in a held-out test set used for the final evaluation, we obtained equal performance. As shown in Table 1, to benchmark our model, we used an extensive set of machine learning classifiers to compare the model performance. Overall, EpiBERTope performs higher in all metrics than other ML algorithms, which are all rigorously hyper-parameter tuned from 5-fold cross-validation.

**Table 1.** Benchmark results of the linear epitope prediction task. Bold texts represent state-of-the-art results.

| Algorithm | AUC | Macro F-1 | Micro F-1 | Precision | Recall |
| --- | --- | --- | --- | --- | --- |
| EpiBERTope | **0.922** | **0.853** | **0.860** | **0.855** | **0.852** |
| EpiBERTope with TAPE pretrain | 0.685 | 0.634 | 0.635 | 0.648 | 0.652 |
| EpiBERTope without pretrain | 0.557 | 0.509 | 0.509 | 0.538 | 0.537 |
| BaggingClassifier | 0.861 | 0.752 | 0.771 | 0.747 | 0.767 |
| RandomForest Classifier | 0.857 | 0.762 | 0.780 | 0.756 | 0.780 |
| ExtraTrees Classifier | 0.857 | 0.763 | 0.781 | 0.756 | 0.780 |
| GradientBoosting Classifier | 0.736 | 0.661 | 0.699 | 0.659 | 0.695 |
| AdaBoost Classifier | 0.590 | 0.489 | 0.611 | 0.535 | 0.586 |
| LogisticRegression Classifier | 0.590 | 0.588 | 0.611 | 0.535 | 0.568 |
| LinearDiscriminant Analysis | 0.590 | 0.490 | 0.612 | 0.536 | 0.586 |
| BernoulliNB | 0.588 | 0.518 | 0.609 | 0.544 | 0.576 |
| GaussianNB | 0.583 | 0.536 | 0.536 | 0.558 | 0.558 |
| Quadratic Discriminant Analysis | 0.515 | 0.506 | 0.549 | 0.511 | 0.512 |
| SupportVector Classifier | 0.510 | 0.384 | 0.601 | 0.502 | 0.570 |

In the linear epitope prediction task, the dataset comprises a total of 11,834 epitopes and 17,722 non-epitopes(See Methods 2.1.2), enabling the models to capture features of linear epitopes from a uniform size of epitope dataset. Here, tree-based algorithms are found to be more effective than boosting-based methods in modeling epitope sequences.

### 3.2 Swiss-Prot pre-training understands protein sequence features efficiently

As shown in Figure 2, the model masks 15% of the sequence tokens to predict which amino acid should be proper in the masked region (See Methods 2.3). This masking strategy enables the model to learn the basic structure and combinations of residues of protein sequences.

To validate the contribution of Swiss-Prot pre-training, we replaced Swiss-Prot with TAPE, a well-established language model specifically designed for modeling proteins. Since TAPE uses a broader collection of protein sequences from the UniProt database that includes the *in-silico* dataset [29], it was difficult for the model to understand sequence context despite the large epitope dataset. The result shows that pre-training on curated proteins (Swiss-Prot) is highly effective in capturing linear epitope sequence information.

We also randomly initialized the pre-training to test if the model can learn only from the epitope dataset. As expected, due to a large number of parameters, the without-pretraining model achieved an AUC of 0.557 (See Table 2).

**Table 2.** Benchmark results of the linear and structural epitope. Bold texts represent state-of-the-art results.

| Algorithm | AUC | Macro F-1 | Micro F-1 | Precision | Recall |
| --- | --- | --- | --- | --- | --- |
| *Structural epitope* |  |  |  |  |  |
| EpiBERTope | **0.677** | **0.524** | 0.731 | **0.539** | **0.605** |
| EpiBERTope with TAPE pretrain | 0.616 | 0.513 | **0.747** | 0.526 | 0.563 |
| EpiBERTope without pretrain | 0.564 | 0.469 | 0.661 | 0.511 | 0.532 |
| Bepipred-2.0 | 0.62 | - | - | - | - |
| LBtope | 0.54 | - | - | - | - |
| NetSurfP | 0.60 | - | - | - | - |
| *Linear epitope* |  |  |  |  |  |
| EpiBERTope | **0.587** | 0.503 | **0.608** | **0.576** | 0.538 |
| EpiBERTope with TAPE pretrain | 0.561 | **0.525** | 0.593 | 0.555 | **0.539** |
| EpiBERTope without pretrain | 0.521 | 0.514 | 0.531 | 0.514 | 0.514 |
| Bepipred-2.0 | 0.574 | - | - | - | - |

### 3.3 EpiBERTope predicts both structural and linear epitopes with high accuracy

We also validated the model’s performance on predicting based on the structural epitope dataset. Although the experimental assay used to obtain a structural dataset considers the structure of the protein sequence, we believe that the sequence-based EpiBERTope model can capture long-distance interactions without introducing structure information of the sequences. As proof of concept, we fine-tuned EpiBERTope on the structural epitope dataset and predicted on left-out linear and structural epitope datasets. For model validation, we used a total of 6 antigen sequences for the structural dataset (See Methods 2.1.3) and the entire linear epitope dataset (See Methods 2.1.2).

As shown in Table 2, EpiBERTope outperforms in AUC for two of the modified EpiBERTope models that are pre-trained with TAPE and without pre-training respectively for both linear and structural epitope datasets. Likewise, the model outperforms a random forest-based Bepipred-2.0 model [11], a linear-based model called LBtope [30], and a baseline predictor that uses RSA values from NetSurfP [31]. This result is promising that the sequenced-based attention mechanism (Equation 3) captures long-range interactions in protein 3D structure more effectively than the models with an additional set of features including computed volume, polarity, calculated surface accessibility (RSA), and secondary structure of each residue.

The model performance is relatively low due to the limited size of the training dataset (total of 770 sequences) compared to training with the linear epitope dataset. Furthermore, we suspect that predicting linear epitopes from the model trained on structural epitopes was a challenging task due to the different modalities of the two datasets, leading to poor performance compared to other tasks.

## 4 Discussion

The Epitope prediction task has been a long-standing challenge, and many methods have been developed to solve this problem [32]. These methods rely on machine learning classifiers by learning features that best differentiate epitopes against non-epitopes. Such methods include SEPPA (a logistic regression model that aims to maximize likelihood function; [33]), BEST (support vector machine that finds hyperplane in N-dimensional space that best classifies data points; [34]), and BepiPred 2.0 (Random forest model that aggregates outputs of individual decision trees, reducing the variance from single decision tree; [11]). These methods do not consider interactions between amino acids, which are central for the stabilization of protein structure and complexes [35].

To overcome the limitation of traditional ML models, we developed a transformer-based model called EpiBERTope that utilizes a multi-head attention mechanism to construct global dependencies of each amino acid in protein sequences. The model is divided into two modules: pre-training and fine-tuning. The pre-training module is a masked language model that explores the grammar of protein sequences by randomly masking the subset of the amino acids and predicting the masked regions. The fine-tuning stage is specifically tailored to the epitope classification task, where we feed in both linear and structural epitope datasets to model the sequences.

Intriguingly, learning global dependencies of the sequence through attention-mechanism is found to be effective in learning epitope properties. We demonstrated that EpiBERTope outperforms highly optimized machine learning models including random forest and SVM on linear epitope prediction tasks. To obtain a more generalizable model across different datasets, we applied the same model to the structural epitope dataset. Here EpiBERTope consistently outperformed, suggesting the task-independent capability of the model.

Moreover, we demonstrated that the pre-training part of EpiBERTope is an essential component of the model. Following the same fine-tuning procedure, we (i) substituted pre-training embedding with a robust TAPE pre-trained model [19] and (ii) randomly initialized the weights in the pre-training model to remove the effect. In both cases, EpiBERTope with curated Swiss-Prot pre-training dataset was the most effective learning sequence properties to identify epitopes. Nevertheless, the model faces one of the current limitations, which is the heterogeneity of antigen sequences and scarcity of structural dataset. For epitope prediction using structural dataset as pre-training, the model suffers from the limited size of the structural dataset and struggles to explore vast domains of protein sequences. Furthermore, predicting linear epitopes from the learned model on structural epitopes was even more challenging due to the heterogeneous distribution of the two data types.

Our approach to predicting epitope is solely based on antigen sequences. Other language models that integrate additional features of the protein sequence such as domain structures, post-translational modifications, or variants [23] can lead to better performance. In general, more complexity introduced to the model has greater potential to capture undiscovered relationships between amino acids.

For future works, EpiBERTope has numerous potential applications. Essentially, the model can be fine-tuned on any task that requires an understanding of interactions within a sequence. For example, the model can be applied to the T-cell epitope prediction task [36]. Moving on to the antibody’s side of the binding, one can use this model to predict paratopes, which is a critical factor in antibody drug discovery [37]. From a genomic perspective, the model can predict disease variants by understanding the pathogenicity of protein variants [38] or gene expression prediction from a sequence [38], [39]. In conclusion, our model will be an easily applicable yet powerful tool not only for B-cell epitope prediction tasks but also for sequence-related biological problems.

